# PhANNs, a fast and accurate tool and web server to classify phage structural proteins

**DOI:** 10.1101/2020.04.03.023523

**Authors:** Vito Adrian Cantu, Peter Salamon, Victor Seguritan, Jackson Redfield, David Salamon, Robert A. Edwards, Anca M. Segall

## Abstract

For any given bacteriophage genome or phage sequences in metagenomic data sets, we are unable to assign a function to 50-90% of genes. Structural protein-encoding genes constitute a large fraction of the average phage genome and are among the most divergent and difficult-to-identify genes using homology-based methods. To understand the functions encoded by phages, their contributions to their environments, and to help gauge their utility as potential phage therapy agents, we have developed a new approach to classify phage ORFs into ten major classes of structural proteins or into an “other” category. The resulting tool is named PhANNs (Phage Artificial Neural Networks). We built a database of 538,213 manually curated phage protein sequences that we split into eleven subsets (10 for cross-validation, one for testing) using a novel clustering method that ensures there are no homologous proteins between sets yet maintains the maximum sequence diversity for training. An Artificial Neural Network ensemble trained on features extracted from those sets reached a test F_1_-score of 0.875 and test accuracy of 86.2%. PhANNs can rapidly classify proteins into one of the ten classes, and non-phage proteins are classified as “other”, providing a new approach for functional annotation of phage proteins. PhANNs is open source and can be run from our web server or installed locally.

**Author Summary:** Bacteriophages (phages, viruses that infect bacteria) are the most abundant biological entity on Earth. They outnumber bacteria by a factor of ten. As phages are very different within them and from bacteria, and we have comparatively few phage genes in our database, we are unable to assign function to 50%-90% of phage genes. In this work, we developed PhANNs, a machine learning tool that can classify a phage gene as one of ten structural roles, or “other”. This approach does not require a similar gene to be known.

## Introduction

Bacteriophages (phages) are the most abundant biological entity on the Earth (1). They modulate microbial communities by lysing specific components of microbiomes. Via transduction and/or lysogeny, they mediate horizontal transfer of genetic material such as virulence factors (2), metabolic auxiliary genes (3), photosystems and other genes to enhance photosynthesis(4), and phage production in general, by providing the host with immunity from killing by other phages. Temperate phages can become part of the host genome as prophages; most bacterial genomes contain at least one, and often multiple, prophages (5,6).

Phage structures (virions) are composed of proteins that encapsulate and protect their genomes. The structural proteins (or virion proteins) also recognize the host, bind to it and deliver the phage’s genome into the host’s cell. Phage proteins, especially structural ones, vary widely between phages and phage groups, so much so that sequence identity-based methods to assign gene function fail frequently: we are currently unable to assign function to 50-90% of phage genes(7). Experimental methods such as protein sequencing, mass spectrometry, electron microscopy, or crystallography, in conjunction with antibodies against individual proteins, can be used to identify structural proteins but are expensive and time-consuming. A fast and easy-to-use computational approach to predict and classify phage structural proteins would be highly advantageous as part of pipelines for identifying functional roles of proteins of bacteriophage origins. The current increased interest in using phages as therapeutic agents (8,9) demands annotations for as high as possible a fraction of each phage genome, even if they are only provisional.

Machine learning has been used to attack similar biological problems. In 2012, Seguritan et al. (10) developed an Artificial Neural Network (ANN) that used normalized amino acid frequencies and the theoretical isoelectric point to classify viral proteins as structural or not structural with 85.6% accuracy. These ANNs were trained with proteins of viruses from all three domains of life. They also trained two distinct ANNs to classify phage capsid versus phage non-capsid and phage “tail associated” versus phage “non-tail-associated” proteins. Subsequently, several groups have used different machine learning approaches to improve the accuracy of predictions. The resulting tools are summarized in **Table 1.**

**Table 1.** Summary of previous ML-based methods for classifying viral structural proteins.

| Reference | Method | Target proteins | Database size | Accuracy |
| --- | --- | --- | --- | --- |
| Seguritan et al. (10) | ANN | structural (all viruses) versus non-structural (all viruses) | 6,303 structural<br>7,500 non-structural | 85.6% |
| Seguritan et al. (10) | ANN | capsid versus non-capsid (phages only) | 757 capsid<br>10,929 non-capsid | 91.3% |
| Seguritan et al. (10) | ANN | Tail-associated versus non-tail (phages only) | 2,174 tail<br>16,881 non-tail | 79.9% |
| Feng et al.(32) | Naïve Bayes | structural versus non-structural | 99 structural<br>208 non-structural | 79.15% |
| Zhang et al.(33) | Ensemble Random Forest | structural versus non-structural | 253 structural<br>248 non-structural | 85.0% |
| Galiez et al.(11) | SVM | capsid versus non-capsid | 3,888 capsid<br>4,071 non-capsid | 96.8% |
| Galiez et al.(11) | SVM | tail versus non-tail | 2,574 tail<br>4,095 non-tail | 89.4% |
| Manavalan et al. (34) | SVM | structural versus non-structural | 129 structural<br>272 non-structural | 87.0% |
| This work | ANN | Ten distinct phage structural classes plus “others” | 168,660 structural<br>369,553 non-structural | 86.2% |

Each of these previous approaches has important limitations: 1) The classification is limited to two classes of proteins (e.g., “capsid” or “not capsid”). 2) Training and testing sets were small (only a few hundred proteins in some cases), limiting the utility of the approaches beyond those proteins used in testing. 3) Methods that rely on predicting secondary structure (e.g., ViralPro(11)) are slow to run. In general, these newer methods have improved accuracy at the cost of lengthening the time required for training, or have used very small training and/or test sets.

Artificial Neural Networks (ANN) are proven universal approximators of functions in IR^n^,(12) including the mathematical function that maps features extracted from a phage protein sequence to its structural class. We have constructed a manually-curated database of phage structural proteins and have used it to train a feed-forward ANN to assign any phage protein to one of eleven classes (ten structural classes plus a catch-all class labeled “others”). Furthermore, we developed a web server where protein sequences can be uploaded for classification. The full database, as well as the code for PhANNs and the webserver, are available for download at http://edwards.sdsu.edu/phanns and https://github.com/Adrian-Cantu/PhANNs

## Methods

### Database

In this work, we generated two complementary protein databases, “classes” and “others”. The “classes” database contains curated sequences of ten phage structural roles (Major capsid, Minor capsid, Baseplate, Major tail, Minor tail, Portal, Tail fiber, Tail shaft, Collar, and Head-Tail joining). The “others” database contains all phage ORFs that do not encode proteins annotated as “structural” or as any of the 10 categories above.

#### The database of “classes”

Sequences from the ten structural classes were downloaded from NCBI’s protein database using a custom search for the class title (the queries are in the “ncbi_get_structural.py” script in the GitHub repository). We manually removed all sequences whose description didn’t fit the corresponding class.

#### The “others” database

To generate a database for the “others” class, all available phage genomes (8,238) were downloaded from GenBank on 4/13/19. ORFs were found using the GenBank PATRIC(13) server with the phage recipe(14). Sequences annotated as structural or any of the ten classes were removed during manual curation. Furthermore, the remaining sequences were de-replicated at 60% together with sequences in the “classes” database using CD-hit(15). Any phage ORF that clustered with a sequence from the “classes” database was removed from the “others” database.

#### Training, test, and validation split

Each class was clustered at 40% using CD-hit and split into eleven sets (10 for cross validation and one for testing, as shown in **Figure 1**). Once the clusters were determined, to prevent loss of the sequence diversity available within the clusters, which is essential for optimal training, the clusters were expanded by adding back *within* each set all the representatives of that set (described in Figure 1). Subsequently, the sets corresponding to each structural class were merged. We named the generated sets 1D-10D and TEST. Splitting the database this way ensures that the different sets share no homologous proteins, yet recapture all the sequence diversity present in each class. Finally, 100% dereplication was performed to remove identical sequences (See **Table 2**). The effect of the cluster expansion on performance is explored in **Figures S1 and S2**.

**Figure 1.**
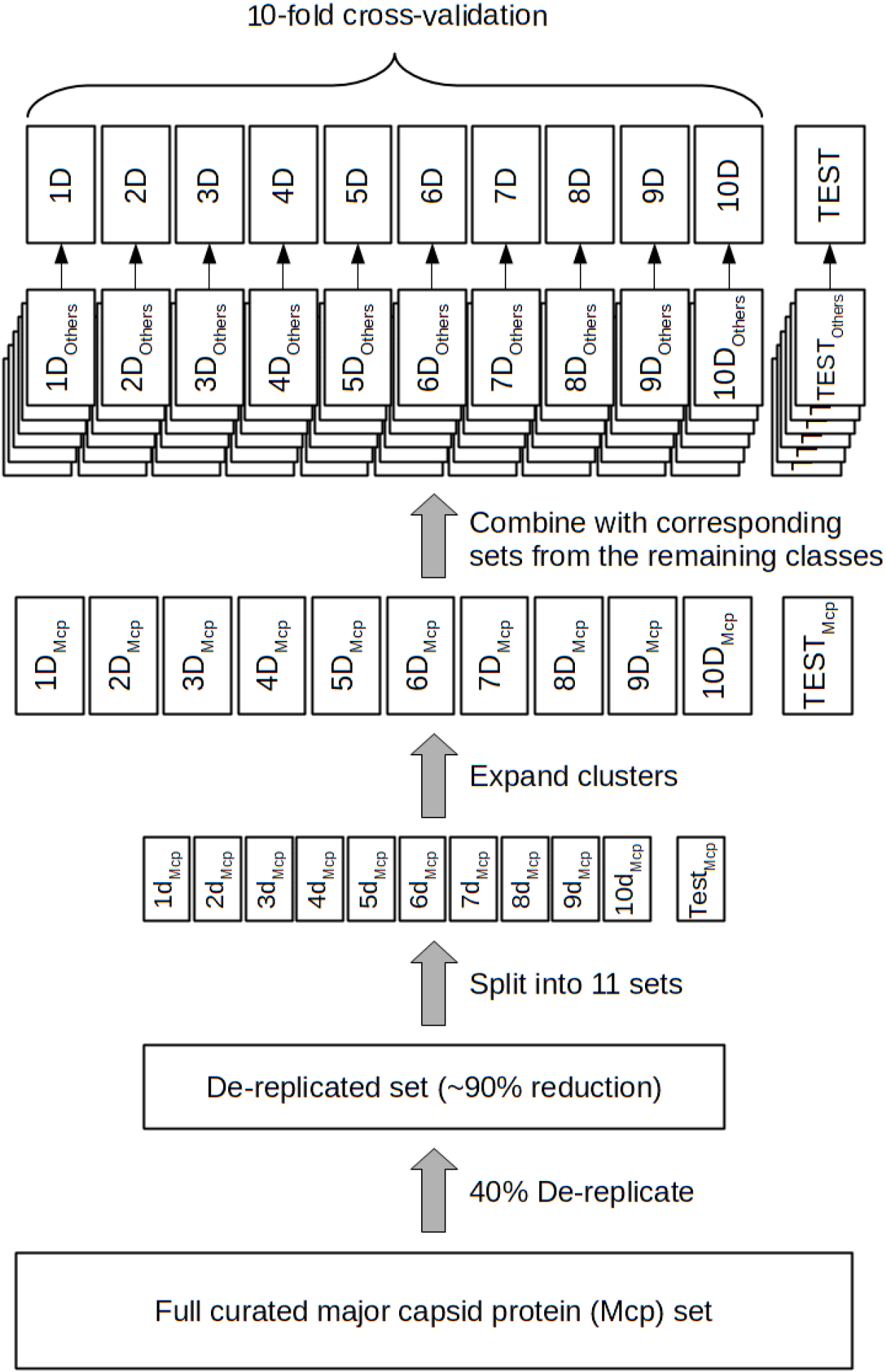
Non homologous database split. To ensure that no homologous sequences are shared between the test, validation, and training sets the sequences from each class (Major capsid proteins in this figure) were de-replicated at 40%. In the de-replicated set, no two proteins have more than 40% identity and each sequence is a representative of a larger cluster of related proteins. The de-replicated set is then randomly partitioned into eleven equal size subsets, (1d_Mcp_-10d_Mpc_ plus Test_Mpc_). Those subsets are expanded by replacing each sequence with all the sequences in the cluster it represents (subsets 1D_Mpc_-10D_Mpc_ plus Test_Mpc_). Analogous subsets are generated for the remaining ten classes and corresponding subsets are combined to generate the subsets used for 10-fold cross-validation and testing (1D-10D and TEST).

**Table 2.** Database numbers. Raw sequences were downloaded using a custom script available at https://github.com/Adrian-Cantu/PhANNs. All datasets can be downloaded from the web server. *Numbers before and after removing sequences at least 60% identical to a protein in the classes database.

| Class | Raw sequences | After manual curation | After de-replication at 40 % | After expansion and de-replication at 100% |
| --- | --- | --- | --- | --- |
| Major capsid | 112,987 | 105,653 | 1,945 | 35,755 |
| Minor capsid | 2,901 | 1,903 | 261 | 1,055 |
| Baseplate | 75,599 | 19,293 | 401 | 6,221 |
| Major tail | 66,513 | 35,030 | 536 | 7,704 |
| Minor tail | 94,628 | 80,467 | 918 | 18,002 |
| Portal | 210,064 | 189,143 | 2,310 | 59,745 |
| Tail fiber | 29,132 | 18,514 | 1,222 | 7,256 |
| Tail shaft | 37,885 | 35,570 | 599 | 15,349 |
| Collar | 4,224 | 3,709 | 339 | 2,105 |
| Head-Tail joining | 60,270 | 58,658 | 1,317 | 15,468 |
| **Total structural** | **694,203** | **547,940** | **9,848** | **168,660** |
| Others | 733,006 | 643,735/643,380* | 106,004 | 369,553 |

### Extraction of features

The frequency of each dipeptide (400 features) and tripeptide (8,000 features) was computed for each ORF sequence in both the “classes” and “others” databases. As a potential time-saving procedure during neural net training while also permitting classification of more diverse sequences, each amino acid was assigned to one of seven distinct “side chain” chemical groups (**Table S1**). The frequency of the “side chain” 2-mers (49 features), 3-mers (343 features), and 4-mer (2,401 features) was also computed. Finally, some extra features, namely isoelectric point, instability index(16) (whether a protein is likely to degrade rapidly), ORF length, aromaticity(17) (relative frequency of aromatic amino acids), molar extinction coefficient (how much light the protein absorbs) using two methods (assuming reduced cysteins or disulfide bonds), hydrophobicity, GRAVY(18) index (average hydropathy) and molecular weight, were computed using Biopython(19). All 11,201 features were extracted from each of 538,213 protein sequences. The complete training data set can be downloaded from the web server (https://edwards.sdsu.edu/phanns).

### ANN architecture and training

We used Keras(20) with the TensorFlow(21) back-end to train eleven distinct ANN models using a different subset of features. We named the models to indicate which feature sets were used in training: composition of 2-mers/dipeptides (di), 3-mers /tripeptides (tri) or 4-mer/tetrapeptide (tetra), or side chain groups (sc) (as shown in **Table S1**), and whether we included the extra features (p) or not. A twelfth ANN model was trained using all the features (**Table S2**).

Each ANN consists of an input layer, two hidden layers of 200 neurons, and an output layer with 11 neurons (one per class). A dropout function with 0.2 probability was inserted between layers to prevent overfitting. ReLU activation (to introduce non-linearity) was used for all layers except the output, where softmax was used. Loss was computed by categorical cross-entropy and the ANN is trained using the “opt” optimizer until 10 epochs see no training loss reduction. The model at the epoch with the lowest validation loss is used. Class weights inversely proportional to the number of sequences in that class were used.

#### 10-fold cross-validation

Sets 1D to 10D (see **Figure 1**) were used to perform 10-fold cross-validation; ten ANNs were trained as described above sequentially using one set as the validation set and the remaining nine as the training set. The results are summarized in **Figures 2, 3, and 4.**

**Figure 2.**
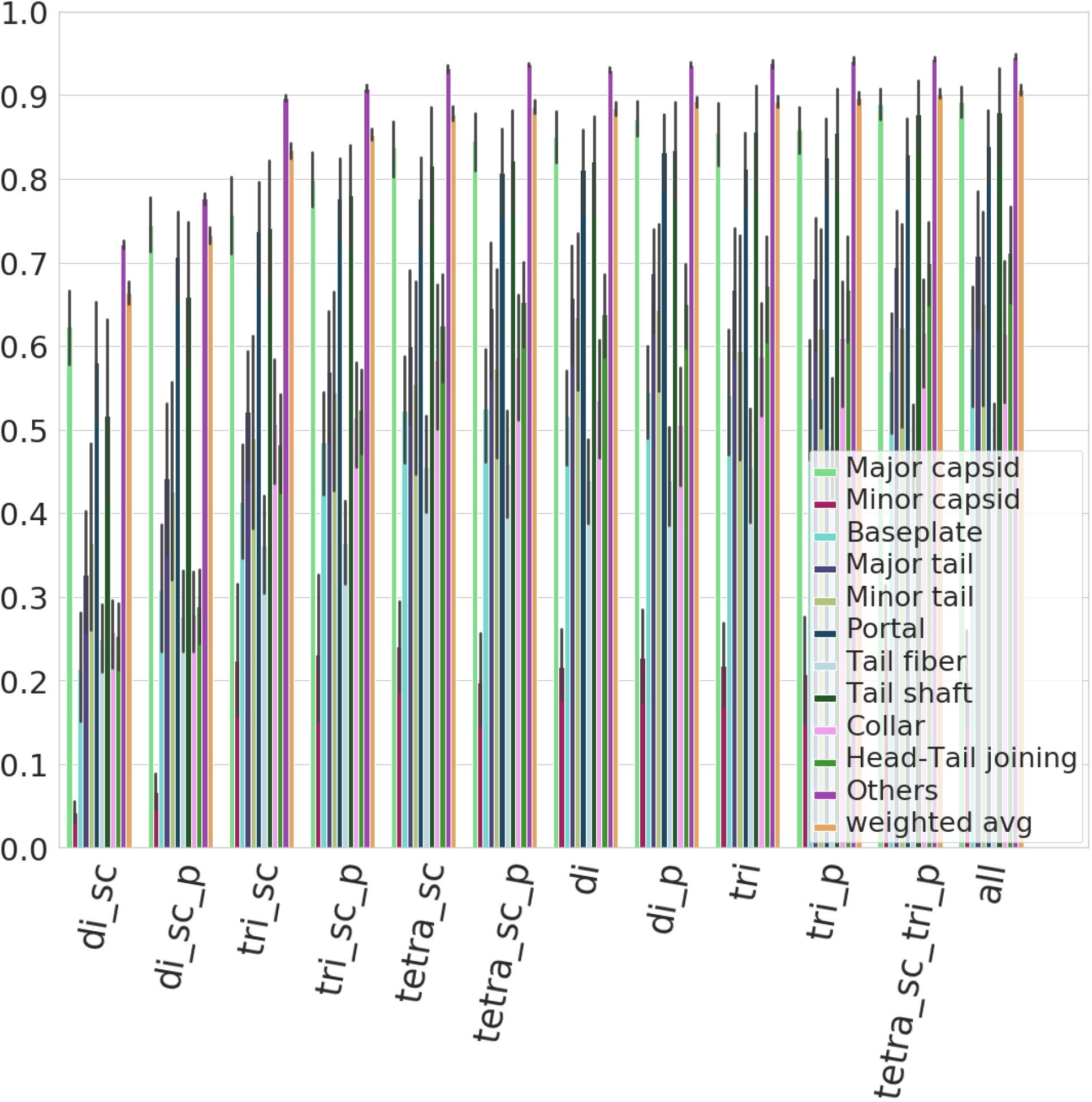
Model-specific F_1_ score. F_1_ scores (harmonic mean of precision and recall) for each model/class combination. All models follow similar trends as to which classes are more or less difficult to classify correctly. Error bars represent the 95% confidence intervals.

**Figure 3.**
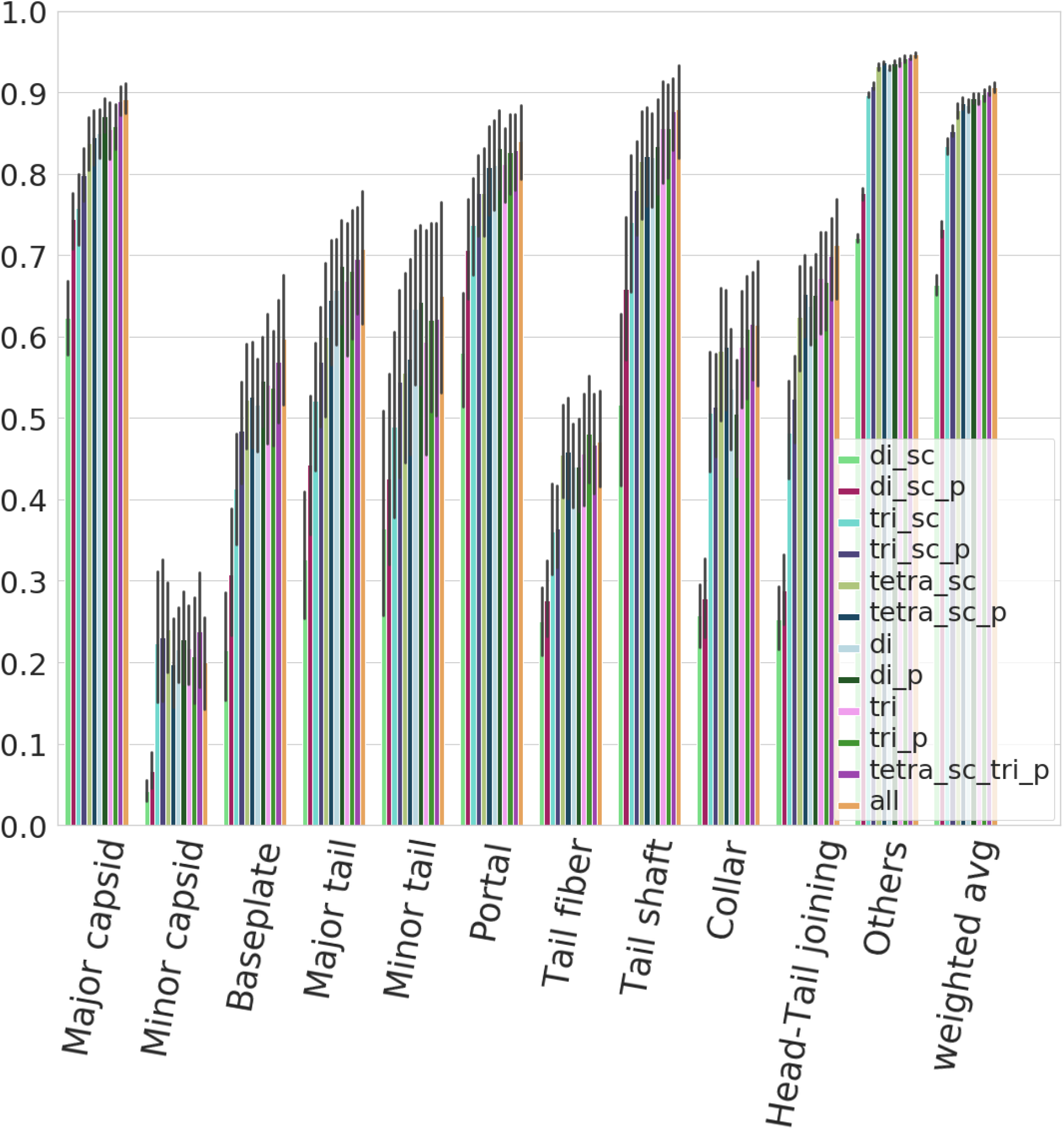
Class-specific F_1_ score. F_1_ scores (harmonic mean of precision and recall) for each model/class combination. Some classes, such as minor capsid, tail fiber, or minor tail, are harder to classify correctly irrespective of the model used. Error bars represent the 95% confidence intervals.

**Figure 4.**
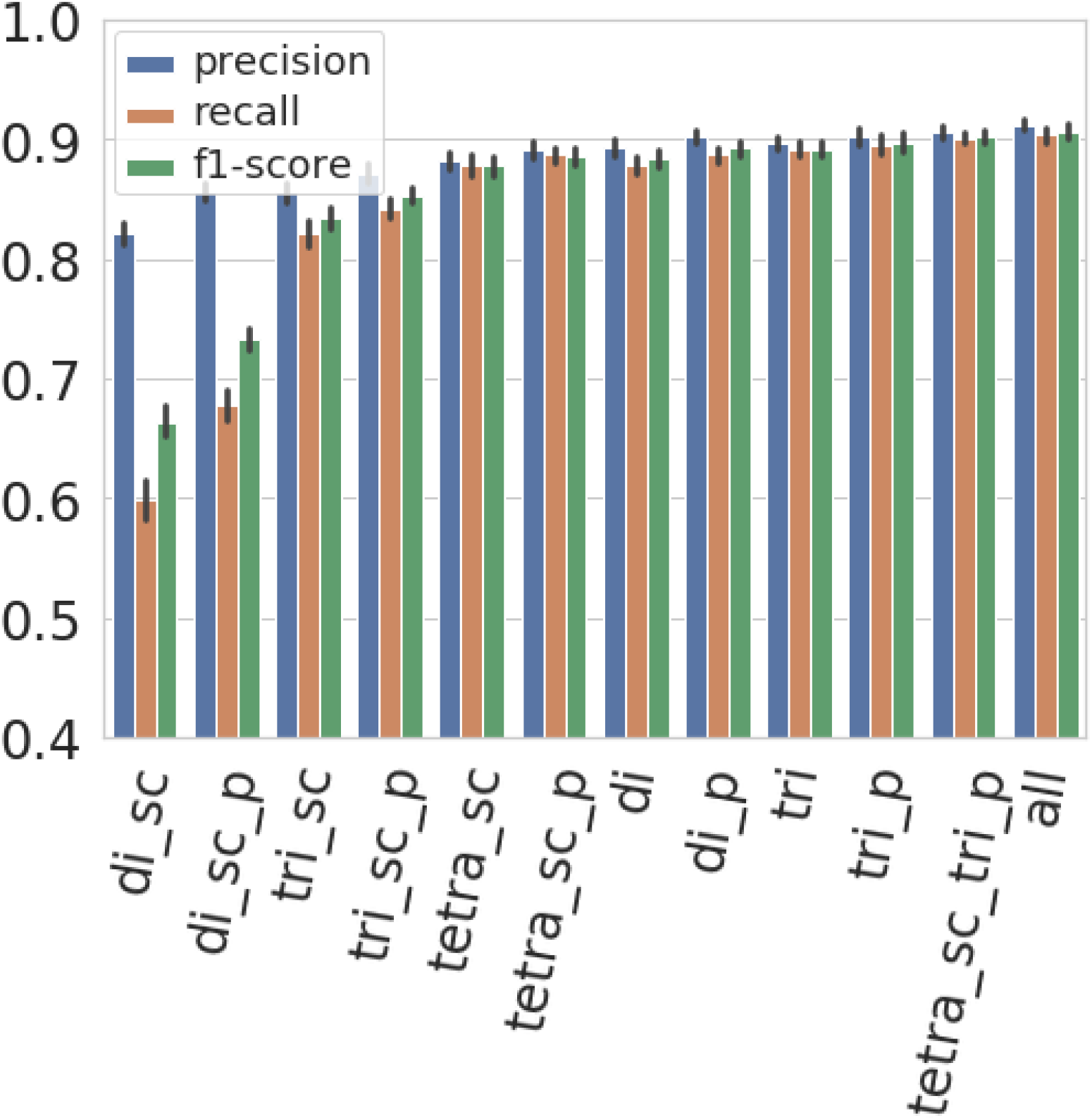
Model-specific weighted average scores. Precision, recall, and F_1_ scores for all Models. Precision is higher in all models as the “others” class is the largest and easiest to classify correctly. Error bars represent the 95% confidence intervals.

### Web server

We developed an easy-to-use web server for users to upload and classify their own sequences. Although ANNs need substantial computational resources for training the model (between 54,861 and 127,756,413 parameters need to be tuned, depending on the model), the trained model can make fast *de novo* predictions. Our web server can predict the structural class of an arbitrary protein sequence in seconds and assign all the ORFs in a phage genome to one of the 11 classes in minutes. The application can also be downloaded and run locally for high-throughput queries or if privacy is a concern.

## Results and discussion

We evaluated the performance of 120 ANNs (10 per model type) on their respective validation set. For each ANN, we computed the precision, recall, and F_1_-score of the 11 classes. A “weighted average” precision, recall and F_1_-score, where the score for each class is weighted by the number of proteins in that class (larger classes contribute more to the score) was computed. The weighted average is represented as a 12th class (**see Table S3**). This gives us ten observations for each combination of model type and class, which allows us to construct confidence intervals as those seen in **Figures 2, 3, and 4.**

**Figure 2** shows that all the models follow the same trend as to which classes they predict with higher or lower accuracy. Some classes of proteins, for example major capsids, collars, and head-tail joining proteins, are predicted with high accuracy. On the other hand, the minor capsid and tail fiber protein classes seem to be intrinsically hard to predict with high accuracy irrespective of the model type used (**Figure 3**). One reason for this is the limited size of the training set: the minor capsid protein set is the smallest class, with only 581 proteins available for inclusion in our database. Even if the classes were weighted according to their size during training, it appears we do not have enough training examples to correctly identify this class. Furthermore, “minor capsid” is often misclassified as “portal” (**Figure 5**). This is probably an annotation bias, as there were about 800 proteins annotated as “portal (minor capsid)” in the raw sequences. When the ~800 proteins are analyzed with PhANNs, over 90% are predicted to be portal proteins. Although these were removed during manual curation of the training data sets, some (small fraction of) minor capsid proteins in our database may have been annotated as “minor capsid” by homology to one of those 800 sequences.

**Figure 5.**
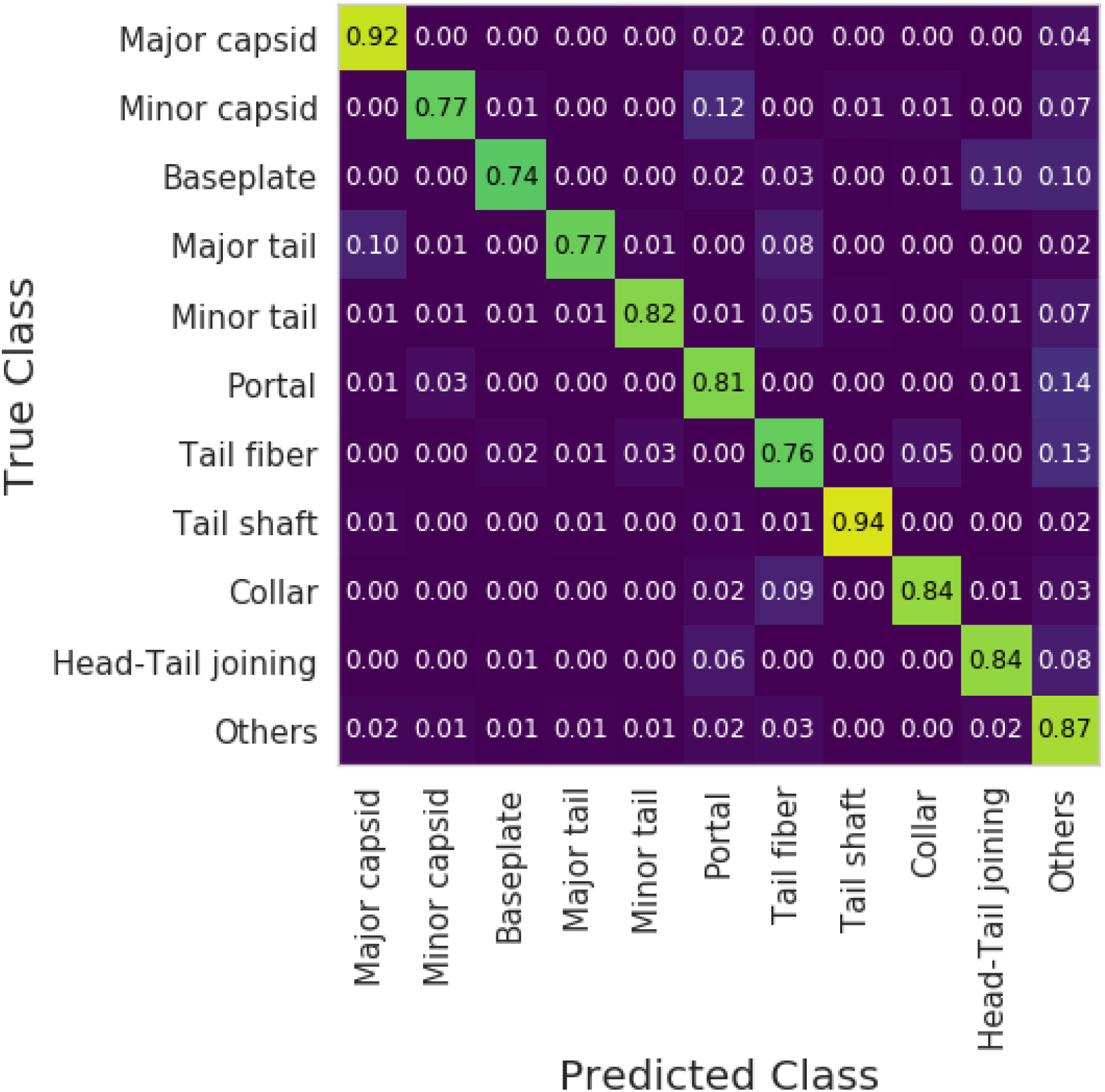
Confusion matrix using the “tetra_sc_tri_p” model. Each row shows the proportional classification of test sequences from a particular class. A perfect classifier would have 1 on the diagonal and 0 elsewhere. In general, a protein that is misclassified is predicted as “others”.

The predictive accuracy for a specific class of proteins is likely to be affected by the bias in the training datasets. The bias could be biological and/or due to a sampling bias. An example of the former is the tail fiber class: the tail fiber is one of the determinants of the host range of the virus, and is under strong evolutionary selective pressure (22–27). On the other hand, sampling bias may be introduced due to oversampling of certain types of phages, such as the thousands of mycobacterial phages isolated as part of the SEA-PHAGES project(28); many of which are highly related to each other.

Average validation F_1_-scores range from 0.664 for the “di_sc” model to 0.906 for the “all” model (**Figure 4**). Although the average validation F_1_-score for the top three models “tri_p” (0.897), “tetra_sc_tri_p” (0.901), and “all” (0.906) are not significantly different from each other, we decided to use “tetra_sc_tri_p” for the web server because, while it uses ~7% fewer features than “all” (10,409 vs 11,201), we expect that the tetra side chain features will be better than the tripeptide features at generalizing predictions and accessing greater sequence diversity.

Using the “tetra_sc_tri_p” ensemble, we predicted the class of each sequence in the test set (47,879) by averaging the scores of each of the ten ANNs. Results are summarized in **Figure 5**. Doing this we reach a test F_1_-score of 0.875 and accuracy of 86.2% over the eleven classes.

The performance of any machine learning system is limited by the availability and cost of training examples (29). This has resulted in top performing image and audio classification systems invariably augmenting their training data with synthetic examples created by applying semantically orthogonal transformations to the training set (slightly rotating or distorting an image, adding background noise to audio) (30,31). In bioinformatics, the current practice of de-replication moves us in exactly the opposite direction -- perfectly good samples cannot be used if their overlap with other samples is too high, leaving only one version of the biostring to learn, and thereby ignoring any variations. Our approach overcomes this failing by using all non-redundant data. By splitting the dataset into the training, validation and test sets after first de-replicating at 40%, we remove even slightly redundant samples and make sure that none of the performance is due to memorization rather than generalization. Augmenting the training set by expanding the clusters back out to all non-redundant samples is the novel idea we have introduced in the present paper as a way of increasing our training set size and hence our accuracy.

## Conclusion

ANNs are a powerful tool to classify phage structural proteins when homology-based alignments do not provide usable functional predictions, such as “hypothetical” or “unknown function”. This approach will get more accurate as more and better characterized phage structural protein sequences, especially more divergent ones, are experimentally validated and become available for inclusion in our training sets. This method can also be applied to predicting the function of unknown proteins of prophage origin in bacterial genomes. In the future, we plan to expand this approach to more protein classes and to viruses of eukaryotes and archaea.

## Acknowledgments

### Funding

This research is based upon work supported by the Office of the Director of National Intelligence (ODNI), Intelligence Advanced Research Projects Activity (IARPA), via the Army Research Office (ARO) under cooperative Agreement Number W911NF-17-2-0105, and awarded as a partial subcontract to AMS. The views and conclusions contained herein are those of the authors and should not be interpreted as necessarily representing the official policies or endorsements, either expressed or implied, of the ODNI, IARPA, ARO, or the U.S. Government. The U.S. Government is authorized to reproduce and distribute reprints for Governmental purposes notwithstanding any copyright annotation thereon.

Victor Seguritan and Jackson Redfield were supported by NSF DMS 0827278 Undergraduate BioMath Education grant awarded to AMS and PS.

---

**Table S1.** Side chain groupings.

|  |  |
| --- | --- |
| Hydrophobic | A,I,L,M,V |
| Hydrophilic | N,Q,S,T |
| Small turn | G,P |
| Disulfide | C |
| Positive charge | H,K,R |
| Negative charge | D,E |
| Aromatic | F,W,Y |

**Table S2.**
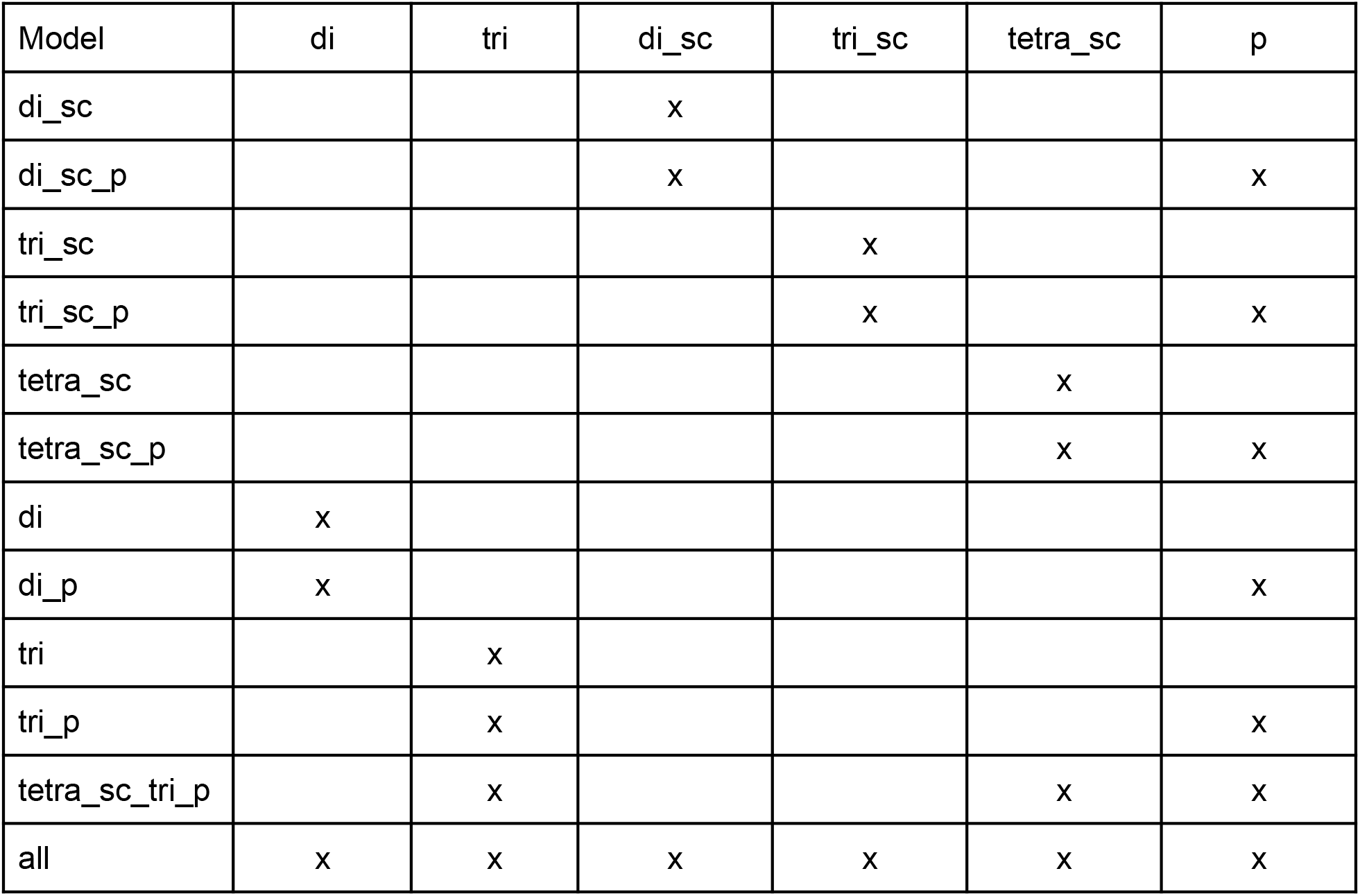
Feature types included in each of the 12 models. **di** - 2-mer/dipeptide composition; **tri** - 3-mer/tripeptide composition; **tetra** - 4-mer/tetrapeptide composition; sc - side-chain grouping; **p** - plus all the extra features [isoelectric point, instability index (whether a protein is likely to be degraded rapidly), ORF length, aromaticity (relative frequency of aromatic amino acids), molar extinction coefficient (how much light a protein absorbs) using two methods (assuming reduced cysteines or disulfide bonds), hydrophobicity, GRAVY index (average hydropathy), and molecular weight, as computed using Biopython].

**Figure S1.**
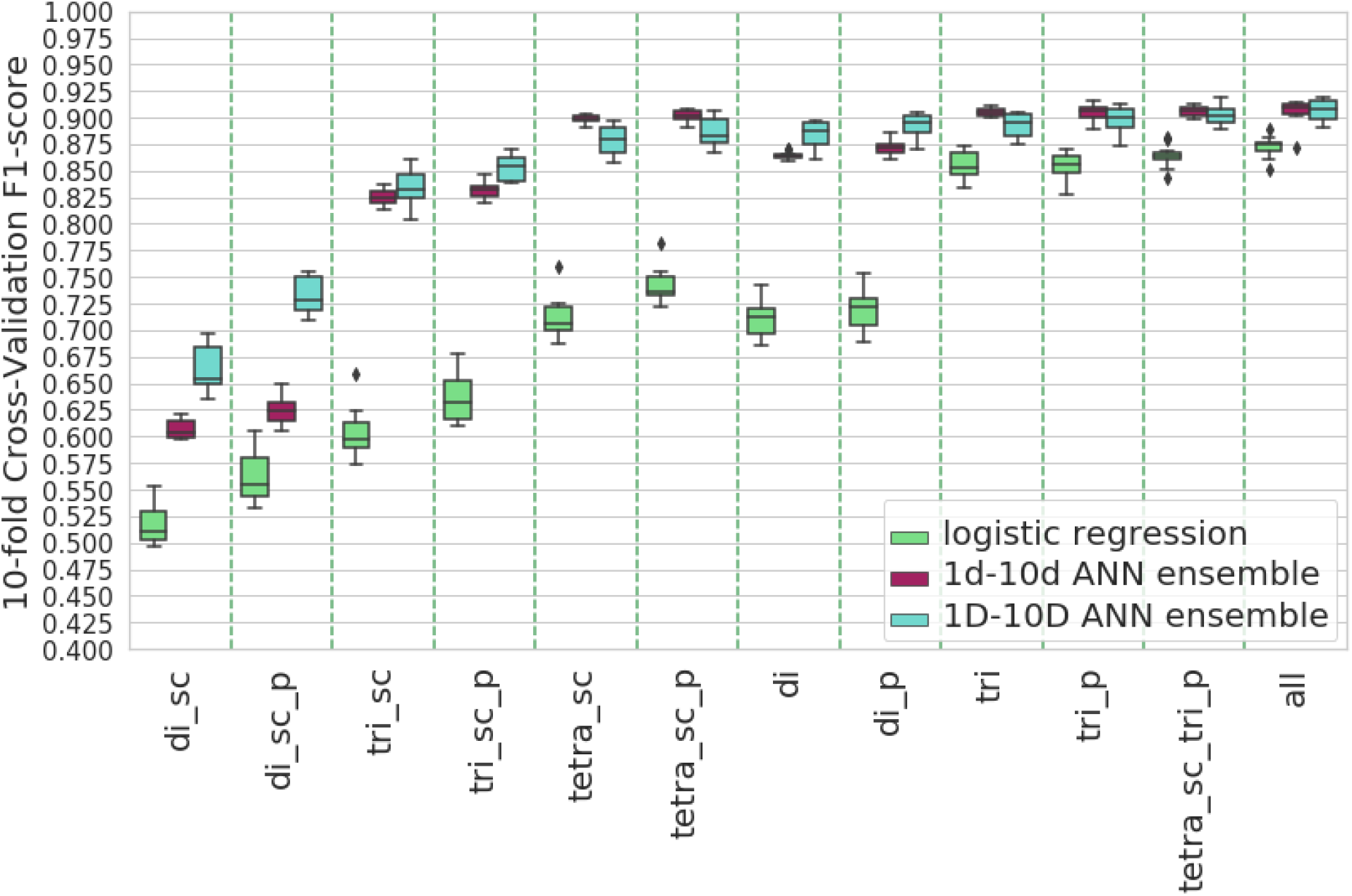
Comparison of the validation weighted average F_1_-score of three models on the same feature sets. We compared our ANN ensemble trained on 1D-10D sets against a logistic regression trained on the 1D-10D sets and an ANN ensemble trained on the 1d-10d sets (40% dereplication, without cluster expansion --see Methods). The ANN ensembles perform significantly better than the logistic regression. Error bars represent 0.95 confidence intervals.

**Figure S2.**
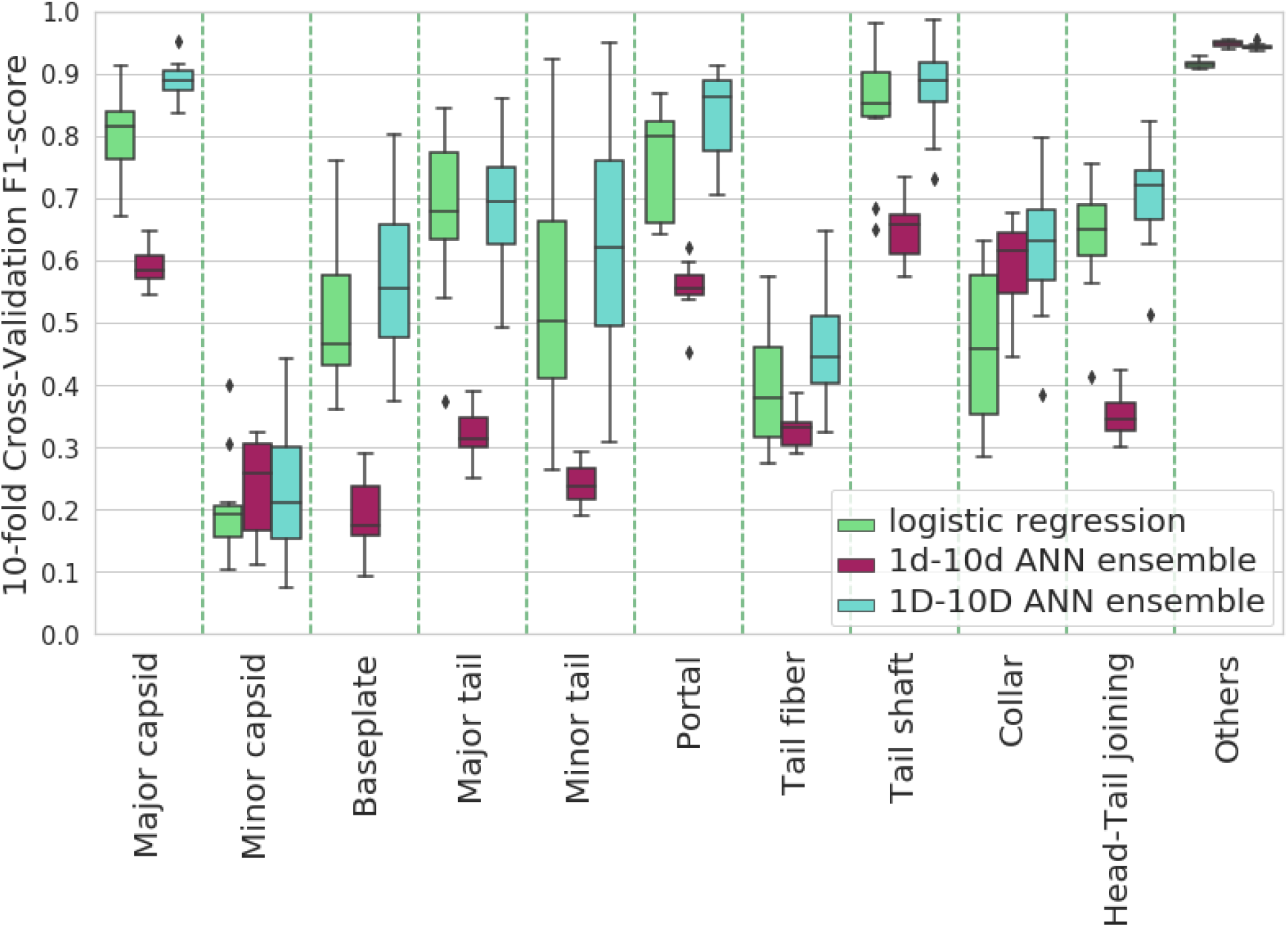
Per class comparison of the validation F_1_-score of three models on the “tetras_sc_tri_p feature” set. In the structural classes, the 1D-q0D ANN ensemble performs slightly better than the logistic regression and significantly better than the 1d-10d ANN ensemble. In the “others” class (by much the largest), 1D-10D ANN ensemble performs as well as 1d-10d ANN and better than logistic regression. Error bars represent 0.95 confidence intervals.

